# Graphics processing unit-accelerated Monte Carlo simulation of polarized light in complex three-dimensional media

**DOI:** 10.1101/2022.01.13.476270

**Authors:** Shijie Yan, Steven L. Jacques, Jessica C. Ramella-Roman, Qianqian Fang

## Abstract

**Significance:** Monte Carlo (MC) methods have been applied for studying interactions between polarized light and biological tissues, but most existing MC codes supporting polarization modeling can only simulate homogeneous or multi-layered domains, resulting in approximations when handling realistic tissue structures.

**Aim:** Over the past decade, the speed of MC simulations has seen dramatic improvement with massively-parallel computing techniques. Developing hardware-accelerated MC simulation algorithms that can accurately model polarized light inside 3-D heterogeneous tissues can greatly expand the utility of polarization in biophotonics applications.

**Approach:** Here we report a highly efficient polarized MC algorithm capable of modeling arbitrarily complex media defined over a voxelated domain. Each voxel of the domain can be associated with spherical scatters of various radii and densities. The Stokes vector of each simulated photon packet is updated through photon propagation, creating spatially resolved polarization measurements over the detectors or domain surface.

**Results:** We have implemented this algorithm in our widely disseminated MC simulator, Monte Carlo eXtreme (MCX). It is validated by comparing with a reference CPU-based simulator in both homogeneous and layered domains, showing excellent agreement and a 931-fold speedup.

**Conclusion:** The polarization-enabled MCX (pMCX) offers biophotonics community an efficient tool to explore polarized light in bio-tissues, and is freely available at http://mcx.space/.

## 1 Introduction

Polarized light has been found to be highly sensitive to medium structures and hence has been widely adopted in optical imaging to probe microstructural features inside biological tissues.^1–4^ For example, the polarization status of the backscattered light can be measured to characterize the superficial layer of skin for cancer diagnostic purposes.^5, 6^ The measurements of tissue birefringence permit quantification of abnormalities of the retinal nerve fiber layer^7^ and cornea,^8^ as well as three-dimensional (3-D) reconstruction of nerve fiber orientations inside human brains^9^ and orientations of collagen within the uterine cervix.^10^ Polarized light imaging (PLI) uses linearly co-polarized images subtracted by those of cross-polarized light to create a differential image based on the small population of superficially scattered photons that still retain much of the incident polarization state.^5^ PLI subtracts the large randomized population of multiple-scattered photons that produce a blinding background of diffuse light. The resulting difference image enhances the contrast of superficial tissue layers and rejects deeper tissue structures, enabling wide-field screening of epidermal or epithelial layers. Mueller Matrix polarimetry and the use of various decomposition methods can also be used to pinpoint different regions and structures within biological tissue.^2, 3^ Accurately simulating polarized light transport inside complex tissues allow quantitative investigations of the depth response of polarized light and the perturbations produced by local tissue abnormalities.

The propagation of polarized light inside scattering media can be described by the vector radiative transfer equation (VRTE).^11^ Analytically solving the VRTE is not viable in complex media such as human tissues. Owing to its high flexibility and simplicity in programming, the Monte Carlo (MC) method, among other numerical techniques,^12–17^ has been one of the limited approaches available to quantitatively model interactions of polarized light with scattering media. Depending on the vectorial representations of polarization states, polarized light MC algorithms can be largely categorized into two formalisms – Jones calculus and Mueller calculus.^2, 4, 18^ The Jones calculus used in the electric-field MC (EMC) algorithm traces the amplitudes and phases of two orthogonal electric field components (Jones vector) and is therefore well suited for simulating light coherence effects.^19^ On the other hand, the Mueller calculus describes the state of polarization using the Stokes vector.^20–24^ The Stokes vector does not contain any phase information but allows to model unpolarized and partially polarized light. The Stokes vector can be obtained by measuring four intensity values. Similarly Mueller matrices can be obtained through sixteen intensity measurements and using Mueller matrix decomposition, quantities such as tissue retardation, depolarization and de-attenuation can be obtained.^4^

A well-known limitation of MC methods is the long computation time. Due to the rapid emergence of massively-parallel computing techniques, benefit largely from the fast advances in many-core processors such as graphics processing units (GPUs), MC simulations of polarized light have seen significant speed improvement over the past decade. Several groups have reported parallel EMC implementations.^25–27^ Wang *et al*.^25^ presented a Compute Unified Device Architecture (CUDA)^28^ based EMC to model coherent light in a single homogeneous slab and achieved over 370 × speedup compared to the CPU counterpart.^19^ Ding *et al*.^27^ extended the GPU-based EMC algorithm to consider multi-layered media at the expense of reduced speedup (~ 45 ×). In addition, Li *et al*.^29^ presented a CUDA-based polarized-light MC algorithm to model interstitial media embedded with spherical and cylindrical scatterers. It employed a single-kernel scheme and was hundreds of times faster than its CPU version.^24^ In 2019, Oulhaj et al.^30^ reported a GPU-accelerated MC algorithm to efficiently compute the sensitivity profile for polarized light inside homogeneous media. The reported GPU implementation was verified against a widely used CPU-based code by Ramella-Roman et al.^22^ and reported over 150× speedup.

Although these studies have demonstrated significantly improved simulation speed, most of these simulators only support layered domains and can not address the needs in modeling increasingly complex media.^4^ In the simulation of biological tissues with irregular-shaped structures, employing simplifications in domain geometries could introduce significant errors. For example, in the MC modeling of human brains, noticeable differences have been observed between layered-slab models and more anatomically realistic models such as voxel-based and mesh-based brain models.^31^

In this work, we present an open-source and GPU-accelerated MC simulator to model polarized light inside 3-D heterogeneous media. This MC algorithm utilizes a 3-D voxelated grid to represent spatially varying distributions of spherical scatterers, characterized by their radii and densities. We use Muller calculus to update the Stokes vectors of simulated photon packets, from which we can compute various polarimetry related measurements along the surface of the domain. In the remainder of this paper, we first briefly review the steps of the meridian-plane MC algorithm.^22^ Then we detail our GPU-implementation of this algorithm, as part of our enhanced open-source MC software – Monte Carlo eXtreme (MCX),^32^ including the preprocessing steps to encode the distribution of particles into a 3-D array data structure and optimization strategies to better use GPU resources. In the Results section, we validate the proposed GPU-based polarization-enabled MCX (pMCX) against the widely used CPU MC simulator “meridianMC” written by Ramella-Roman *et al*.^33^ and quantify the speed improvement using several benchmarks of homogeneous and heterogeneous domains. Finally, we summarize the key findings and discuss future directions.

## 2 Methods

### 2.1 Meridian-Plane Polarized Light MC

The meridian-plane polarized light MC algorithm^33^ largely follows the standard MC photon transport simulation steps,^34^ including “launch”, “move”, “absorb”, “scatter” and “detection”. At the “launch” stage, the initial weight, position and direction vector are defined for each photon packet depending on the profile of the incident beam. To describe the polarization state, the Stokes vector is defined with respect to the initial meridian plane for every simulated photon packet.^22^ The Stokes vector 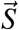 consists of four quantities [*I*, *Q*, *U*, *V*], where *I* (*I* ≥ 0) describes the total light intensity, *Q* (−1 ≤ *Q* ≤ 1) controls the mixing between horizontally (*Q* = 1) and vertically (*Q* = − 1) linearly-polarized light, *U* (−1 ≤ U ≤ 1) controls the mixing between +45° (*U* = 1) and −45° (*U* = −1) linearly polarized light, and *V* (−1 ≤ *V* ≤ 1) controls the mixing between right (*V* = 1) and left (*V* = −1) circularly-polarized light.^1^

After the “launch” step, the photon packet starts propagating inside the simulation domain. In lossy media, the packet weight is monotonically reduced along the photon’s paths and the weight loss is accumulated into the local grid element (such as a voxel or tetrahedral element^35^). When arriving at an interaction site, the photon packet changes direction due to scattering. To compute the new direction cosines, the scattering zenith angle *θ* (0 ≤ *θ* ≤ *π*) and azimuth angle *ϕ* (0 ≤ *ϕ* < 2*π*) are statistically sampled. Compared to the standard MC, the scattering step in a polarized light simulation requires additional computation to properly update 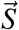. Firstly, the probability density function (also known as the scattering phase function) of polarized light has a bivariate dependence on both *θ* and *φ*. For incident light with a Stokes vector 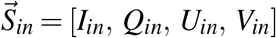, the phase function *P*(*θ*, *φ*) is

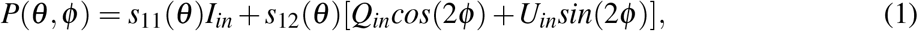

where *s*_11_(*θ*) and *s*_12_(*θ*) are elements from the scattering matrix *M*(*θ*) from a homogeneous spher-ical particle, computed via the Mie theory^36^

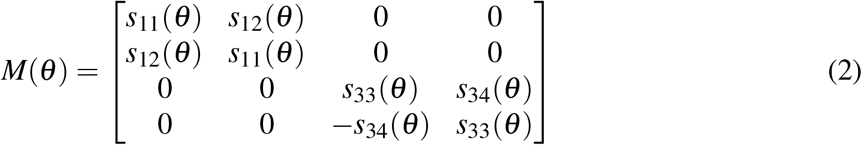

A rejection method^37^ is employed to select angles *θ* and *φ*. Once *θ* and *φ* are determined, the Stokes vector must be rotated relative to the new meridian plane using *M*(*θ*) to update the polarization states.

In order to efficiently perform the rejection method and Stokes vector rotation, the elements of the scattering matrix *M*(*θ*) of all user-specified spherical scatter species are pre-computed over a discreteized set of *θ*. A photon packet is terminated when it escapes from the simulation domain or, if enabled, fails to survive a Russian roulette.^34^ Note that the original meridian-plane polarized light MC assumes refractive-index matched domain boundaries.^22^ The Stokes vector of the escaping photon is rotated relative to the meridian plane of the detector placed immediately outside the domain boundaries, before being accumulated to generate desired output quantities. A detailed description of the formulas used in the meridian plane MC algorithm can be found in the literature.^22^

### 2.2 Implementing Meridian Plane MC in MCX

The original CPU-based meridian-plane MC program,^33^ referred to as “mcMeridian” (`stok1.c`) hereinafter, is dedicated to modeling homogeneous infinite slab geometries. In contrast, the CUDA-based MCX is capable of modeling arbitrarily heterogeneous media represented by a 3-D voxelated domain.^32^ In a non-polarization MCX simulation, the domain is represented by a 3-D integer array with each number representing the index or label of the tissue at each voxel. The actual optical properties of the tissue label are stored in a look-up table, with four element per tissue type: absorption coefficient *μ_a_* (1/mm), scattering coefficient *μ_s_* (1/mm), anisotropy *g* and refractive index *n*. To simulate polarized light, the scattering properties of each type of scatterer must be included in addition. To simply the computation, here we only consider spherical scatterers. The radius *r* (*μ*m), refractive index *n_sph_* and volumetric number density *ρ* (1/*μ*m^3^) of the spherical particle scatters can be specified for each tissue type. When spherical particle properties are specified, the corresponding *μ_s_, g*, and elements of the scattering matrix *M*(*θ*) are pre-computed using the Mie theory^38^ on the host (i.e. CPU). As shown in Eq. 2, the scattering matrix of homogeneous spherical scatterers consists of four independent floating-point numbers *s*_11_(*θ*), *s*_12_(*θ*), *s*_33_(*θ*), and *s*_34_(*θ*). In our implementation, the scattering parameters are sampled at 1,000 evenly spaced points between 0 and *π*, as done in mcMeridian.^33^ The pre-computed optical properties and scattering matrix data are then transferred to the device (i.e. GPU), as illustrated in Fig. 1.

**Fig 1.**
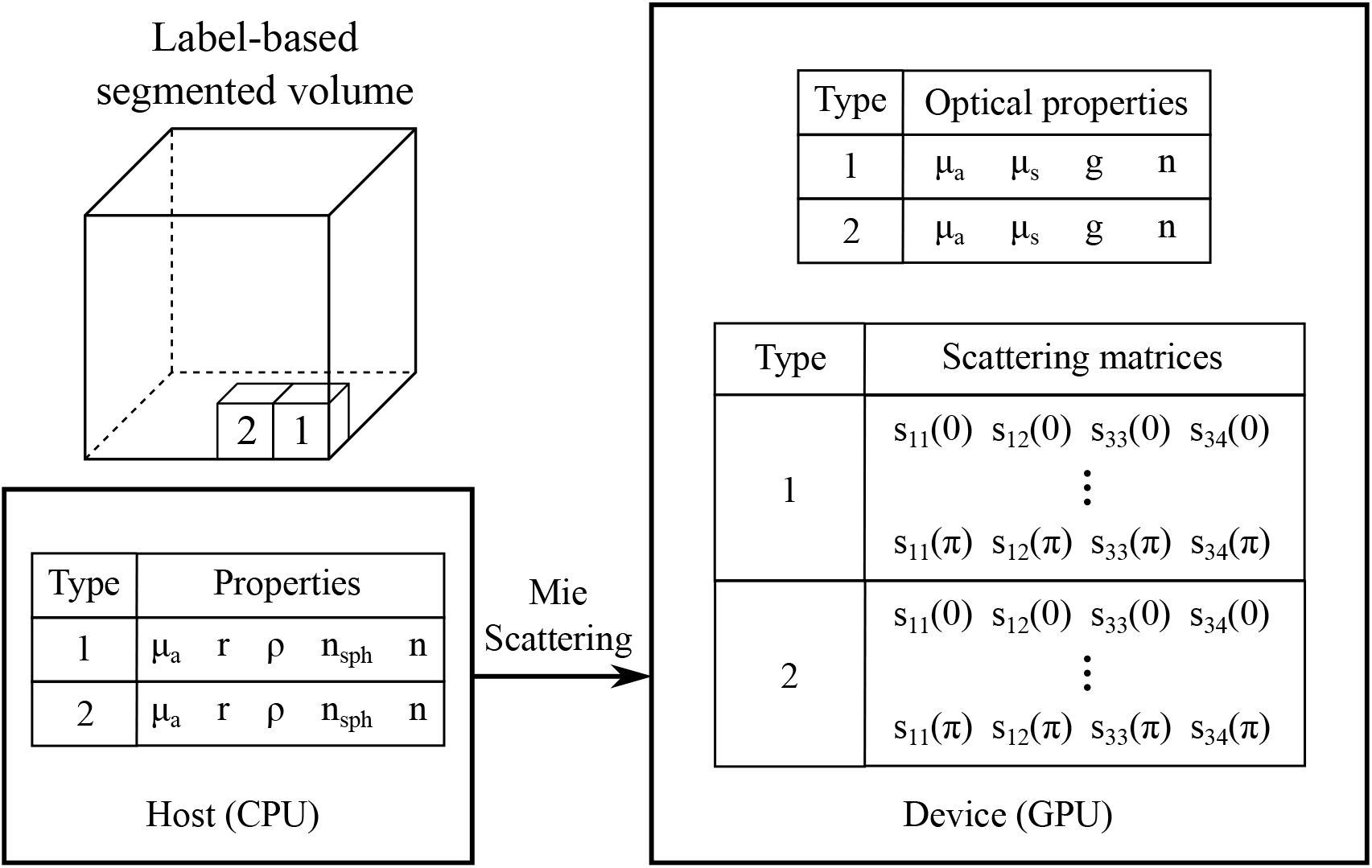
Media representation and media data preprocessing in a polarization-enabled Monte Carlo simulation.

## 3 Results and Discussions

In this section, we first validate the aforementioned polarization-enabled MCX (pMCX) using the single-threaded CPU-based reference implementation^22^ (mcMeridian) in a homogeneous slab. The speed improvement is also quantified. Note that mcMeridian^33^ simulates an infinite slab media geometry in the *x/y* directions while in an MCX simulation, a photon is confined inside a bounding box with user-specified dimensions.^32^ To ensure that our speed comparison is valid, we modified the source code of mcMerdian and added an implicit bounding box to match the dimensions set in pMCX. In the first benchmark, the simulation domain is a 20 × 20 × 10 mm^3^ homogeneous slab, the Mie scattering parameters of the embedded spherical scatterer are *μ_a_* = 0 mm^-1^, *r* = 1.015 μm, *ρ* = 1.152 × 10^-4^ μm^-3^, *n_sph_* = 1.59. The refractive index of the background medium is *n* = 1.33. A monochromatic pencil beam source is positioned at the bottom center (10,10,0) mm of the domain, pointing towards the +*z*-axis and emitting horizontally polarized light at wavelength *λ* = 632.8 nm. The initial Stokes vector of the incident beam is 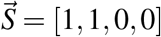. The backscattered photons are collected by a square-shaped area detector (20 × 20 mm^2^) placed on the boundary at *z* = 0 mm. In this benchmark, 10^8^ photon packets are simulated on a desktop running Ubuntu 18.04 with an Intel i7-6700K CPU and an NVIDIA RTX 2080 GPU.

In Fig. 2, we compare the distribution of backscattered [*I*, *Q*, *U*, *V*] components using contour plots in MATLAB (MathWorks, Natick, MA) and observed excellent agreement between mcMeridian and pMCX solutions. We measure the total runtimes, including input data preprocessing, photon transport simulation and output image generation, with mcMeridian reporting 18,111.44 *s* using the Intel CPU and pMCX reporting 19.45 *s* on the NVIDIA RTX 2080 GPU, suggesting a 931 × speedup. In addition, we also benchmark simulation speeds when storing the scattering matrix data over different GPU memory locations, including global, shared and constant memories.^28^ The global memory implementation reports the fastest speed at 8,401 photons/ms, followed by the shared memory (4,965 photons/ms) and constant memory (2,182 photons/ms) implementations. Although the shared memory is known to be the fastest among the three memory types, its has a very small size – up to 48 KB per block.^28^ For storing the scattering matrix of a single species of scatterer at 1,000 angular steps, a total of 16 KB memory is needed. Allocating a large amount of shared memory can lead to drastically reduced active block number, which explains the lower speed compared to the global memory case. On the other hand, constant memory also has a small size (64 KB).^28^ It is most efficient when a memory value is being reused many times after a single read. However, the use of the rejection method requires random access to the buffer which fails to be accelerated by the constant memory due to high “cache-miss”.^28^

**Fig 2.**
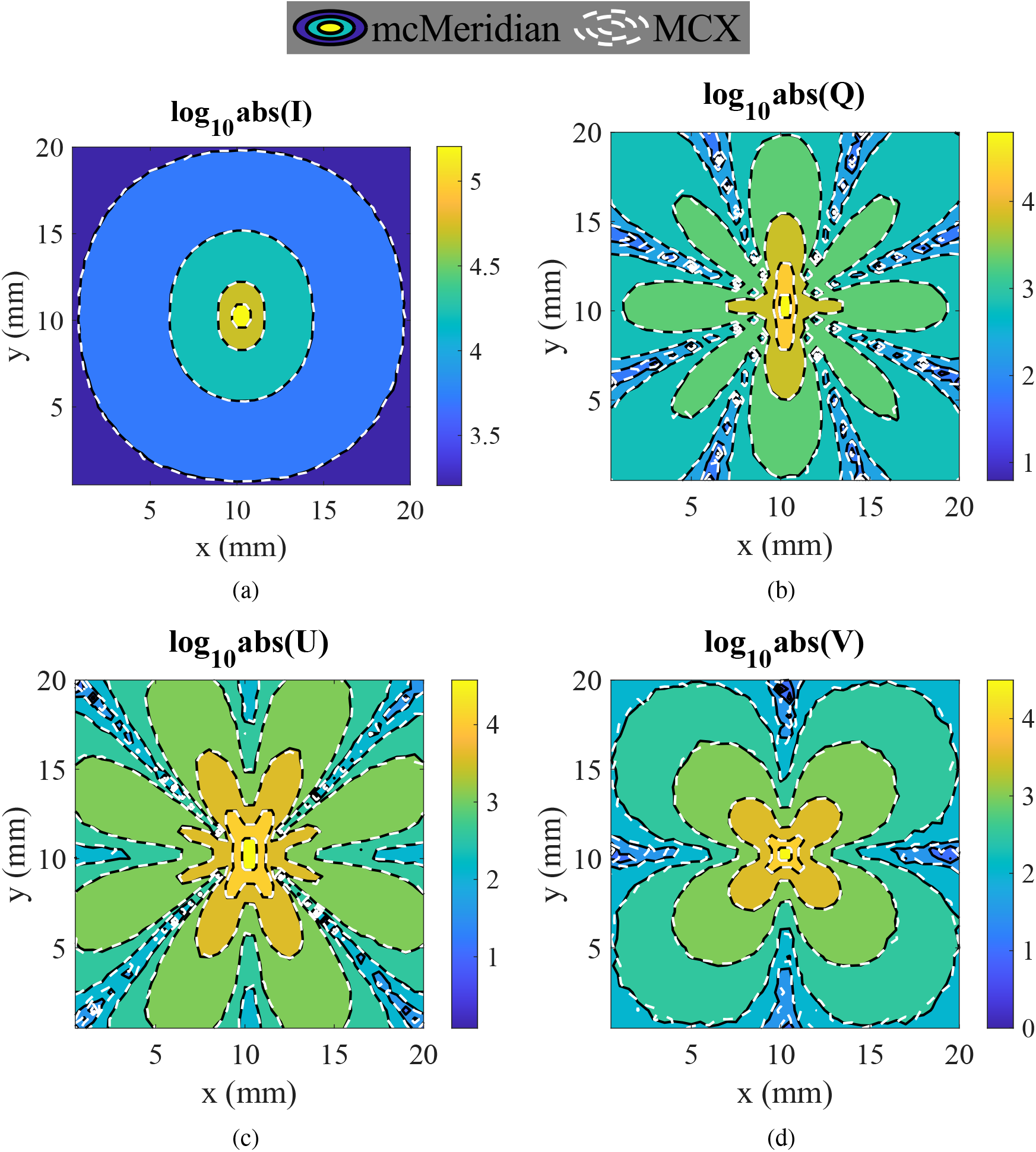
Contour plots of the absolute backscattered [*I*, *Q*, *U*, *V*] (in log_10_ scale) generated by mcMeridian (black solid lines) and pMCX (white dashed lines).

In the next benchmark, we further validate our pMCX simulator by comparing with an extended mcMeridian (with added support of layered media) in a two-layer domain. The slab-shaped simulation domain has a size of 100 × 100 × 50 mm^3^ with the thickness of the superficial layer, *d_e_*, ranging between 0 mm and 10 mm. The Mie scattering parameters are λ = 632.8 nm, *μ_a_* = 0.001 mm^-1^, sphere radius *r* = 0.05 μm, number density *ρ* = 19.11 μm^-3^, *n_sph_* = 1.59, and *n* = 1.33 for the superficial layer. The bottom layer has *r* = 0.3 μm, *ρ* = 2.198 × 10^−2^ μm^-3^, and the other parameters are the same as the superficial layer. These choices of *r* and *ρ* yield a reduced scattering coefficient 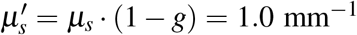 for both layers, along with the same absorption *μ_a_* and hence approximately the same reflected intensity *I* for all *d_e_*. A pencil beam is located on the surface of the slab at (50,50,0) mm, pointing towards the +z-axis and emitting horizontally polarized light. A total of 10^6^ photon packets are simulated for both mcMeridian and pMCX.

In Fig. 3, we plot the total reflected *I* and *Q* components as a function of the superficial layer thickness *d_e_*. The outputs from mcMeridian (black solid lines) and those from pMCX (red circles) once again show excellent agreements. The plot of *Q* increases with the thickness *d_e_* of the superficial layer, which contains smaller spherical scatters and hence stronger back-scattering than the deeper layer, while *I* remains nearly constant as expected. The two-phase transition of *Q* matches our expectations: when the superficial layer is very thin, the reflectance values are close to the value as if the domain is entirely filled with the bottom medium (green dashed line); as we increase *d_e_*, the reflectance values asymptotically approach those determined by the media in the superficial layer (blue dashed line).

**Fig 3.**
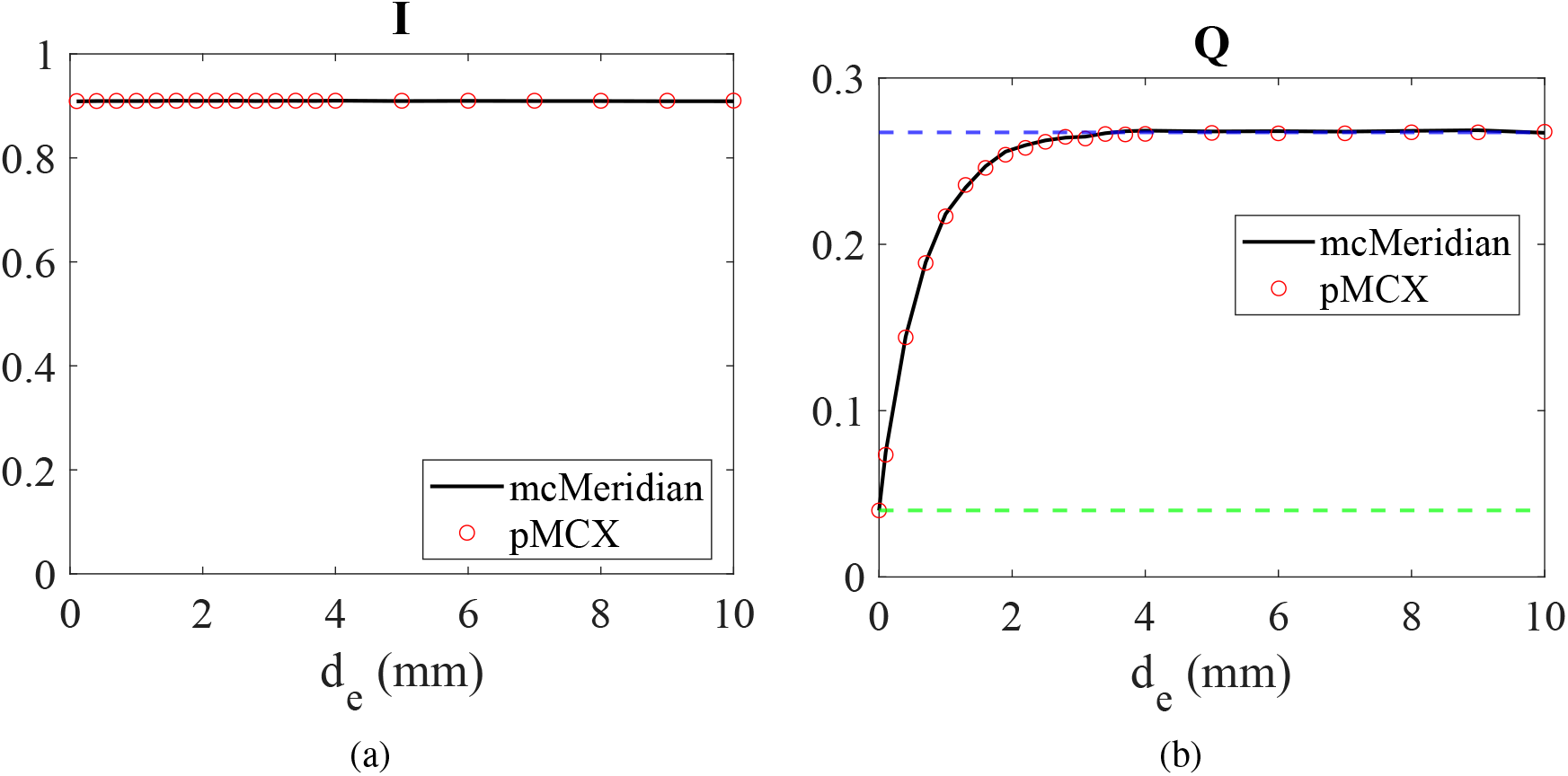
Validation of pMCX in a two-layer domain. We plot the backscattered (a) *I* and (b) *Q* components computed by pMCX and mcMeridian as the superficial layer thickness (d_e_) increases from 0 to 10 mm. Two dashed lines in (b) indicate backscattered *Q* values computed from a homogeneous slab filled only with the medium of the bottom layer (green) and that of the superficial layer (blue).

Finally, we show simulation of a slab-shaped medium embedded with a spherical inclusion, showcasing pMCX’s capability of modeling heterogeneous domains. In this benchmark (Fig. 4), the simulation domain is a 10 × 10 × 1.2 mm^3^ slab with a spherical inclusion of radius 0.5 mm centered at (6,6,0.6) mm. The inclusion and the slab share identical absorption coefficient *μ_a_* = 0.005 mm^-1^, reduced scattering coefficient 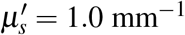, and refractive index *n* = 1.33. However, the Mie scatters inside both domains are different. The background medium is filled with scatterers of radius *r* = 0.05 μm and volume density *ρ* = 19.11 μm^−3^; the inclusion is filled with scatterers of radius *r* = 1 μm and volume density *ρ* = 1.11 × 10^−3^ μm^−3^. The choices of *r* and *ρ* values in either domain was computed based on the Mie theory to ensure their reduced scatter-ing coefficients are the same. All spherical scatterings have a refractive index of, *n_sph_* = 1.59. A 10 × 10 mm^2^ uniform planar light source is placed at the bottom (*z* = 0 mm) surface, pointing towards the +*z*-axis and emitting horizontally polarized light with the wavelength *λ* = 632.8 nm. A cyclic boundary condition is applied to the 4 bounding box facets at ±*x*/±*y* directions to approximate an infinite slab and infinite-plane source. The incident Stokes vector is 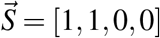. A total of 2 × 10^8^ photons are simulated on an NVIDIA RTX 2080 GPU. We compare the distributions of backscattered [*I*, *Q*, *U*, *V*] at *z* = 0 mm, as shown in Fig. 4. Because the inclusion and background slab share the same *μ_a_*, 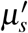 and *n*, a regular diffuse optics forward model without polarization capability would generate no contract to the inclusion. However, our pMCX simulation has revealed distinct image contrasts in *I*, *Q* and *V* images at the correct inclusion locations, suggesting the potential to detect tissue microstructure differences using polarized light.

**Fig 4.**
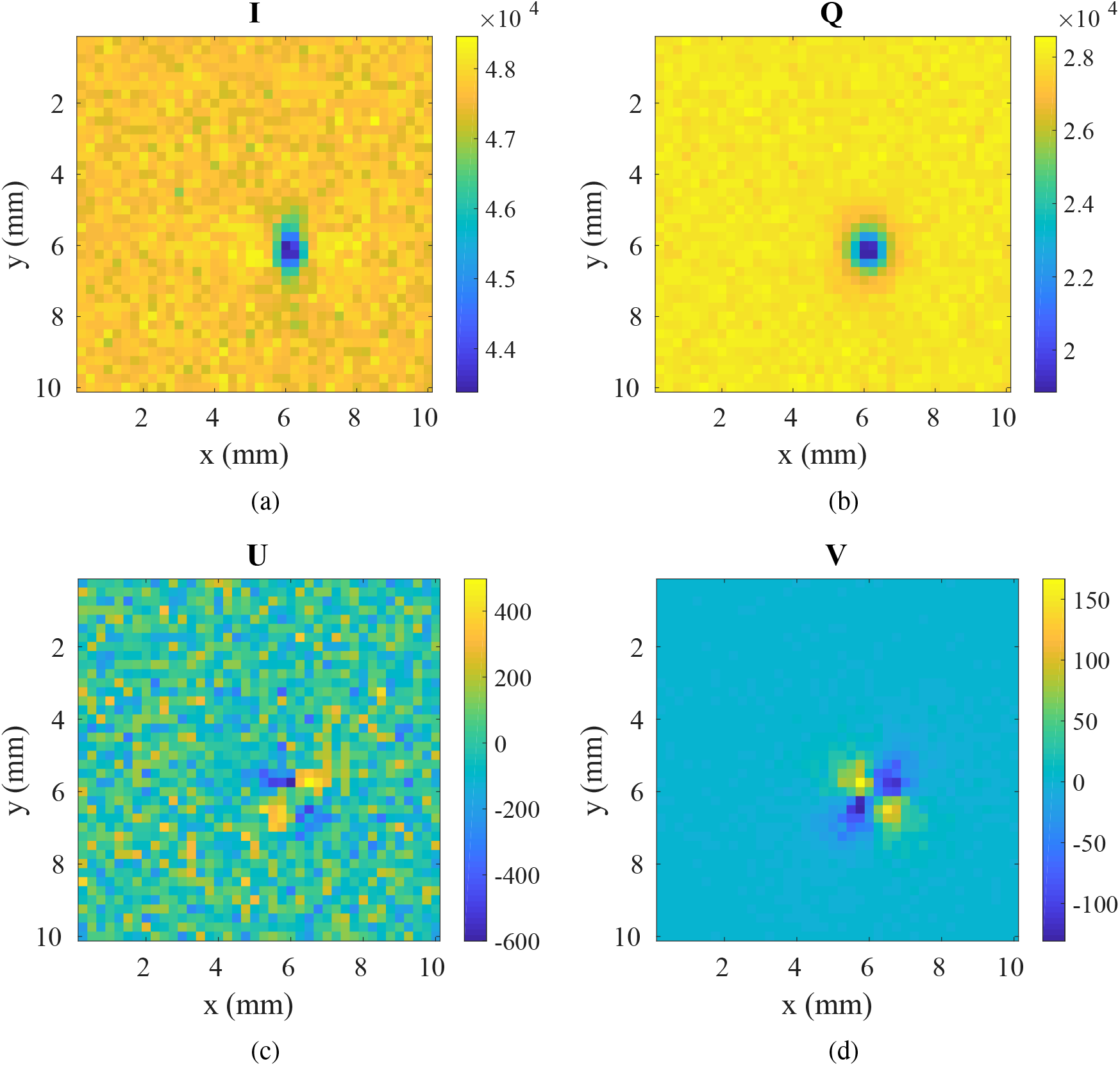
Distributions of [*I*, *Q*, *U*, *V*] backscattered from a10 × 10 × 1.2 mm^3^ slab. A spherical inclusion of radius 0.5 mm is centered at (6,6,0.6) mm.

## 4 Conclusion

In summary, we report a massively-parallel implementation of polarized MC algorithm in our MCX simulator for modeling the propagation of polarized light inside complex media filled with spherical scatterers. Enabled by its built-in voxel-based geometric representation, the polarization-enabled MCX (pMCX) can handle arbitrarily heterogeneous media. We have described the preprocessing steps to encode the scattering properties of spherical particles with various radii and volume densities into a 3-D voxel-based data structure. We provide validation and speed benchmarks ranging from simple homogeneous to complex heterogeneous domains. In all benchmarks, our pMCX solver reports excellent match with the widely used reference solver mcMeridian, while providing a speedup nearly 3 orders of magnitude. In addition, we observed different GPU memory utilization efficiency among global, constant and shared memories, with the global memory implementation yielding the highest speed and least restriction. The pMCX algorithm has been incorporated in our widely disseminated MC simulator and is freely available at http://mcx.space/.

## Acknowledgments

This research is supported by the National Institutes of Health (NIH) grants R01-GM114365, R01-CA204443 and R01-EB026998.

**Shijie Yan** is a PhD student in electrical engineering at Northeastern University. He received his BE degree in information science and engineering from Southeast University, China, in 2013 and MS degree in electrical and computer engineering from Northeastern University in 2017. His research interests include Monte Carlo photon transport simulation algorithms, parallel computing, GPU programming and optimization.

**Steven L. Jacques** is an affiliate professor of bioengineering at the University of Washington in Seattle, Washington. He received his PhD degree from the University of California-Berkeley in 1984, then served as an Research Assoc./Instructor at Massachusetts General Hospital/Harvard Medical School, as an associate professor at the University of Texas M.D. Anderson Cancer Center, and as a full professor at Oregon Health & Science University (OHSU). He has worked in the field of biomedical optics and laser–tissue interactions for 36 years.

**Jessica C. Ramella-Roman** is an Associate Professor in the Bioengineering Department, Florida International University (FIU) in Miami, Florida, USA. She received her PhD degree from Oregon Health & Science University (OHSU) in 2004. She was a postdoctoral fellow at Johns Hopkins University Applied Physics Laboratory and became Assistant Professor at Catholic University of America in 2005, and Associate in 2012 before joining FIU in 2013. Her research focuses on the development of imaging modalities based on spectroscopy and polarization including multimodal applications of nonlinear microscopy.

**Qianqian Fang**, PhD, is currently an Associate Professor in the Bioengineering Department, Northeastern University, Boston, USA. He received his PhD degree from Thayer School of Engineering, Dartmouth College, in 2005. He then joined Massachusetts General Hospital and became an Instructor of Radiology in 2009 and Assistant Professor of Radiology in 2012, before he joined Northeastern University in 2015 as an Assistant Professor. His research interests include translational medical imaging devices, multi-modal imaging, image reconstruction algorithms, and high performance computing tools to facilitate the development of next-generation imaging platforms.

